# Environmentally Relevant Lead Exposure Impacts Gene Expression in SH-SY5Y Cells Throughout Neuronal Differentiation

**DOI:** 10.1101/2024.10.29.620844

**Authors:** Rachel K. Morgan, Anagha Tapaswi, Katelyn M. Polemi, Elizabeth C. Tolrud, Kelly M. Bakulski, Laurie K. Svoboda, Dana C. Dolinoy, Justin A. Colacino

## Abstract

Lead (Pb) causes learning and memory impairments, but the molecular effects of continuous, environmentally relevant levels of exposure on key neurodevelopmental processes are not fully characterized. Here we examine the effects of a range of environmentally relevant Pb concentrations (0.16μM, 1.26μM, and 10μM Pb) relative to control on neural differentiation in the SH-SY5Y cell model. Pb exposure began on differentiation day 5 and was continuous for remaining days, after which we assessed the transcriptome via RNA sequencing at several time points. The bulk of detected changes in gene expression occurred with the 10μM Pb condition. Interestingly, changes associated with the lower two exposures were differentiation stage-specific, with aberrant expression of several genes (e.g., *COL3A1*, *HMOX1*, and *CCL2*) observed during differentiation on days 9, 12, and 15 in both the 0.16μM and 1.26μM Pb conditions, and which disappeared by the time differentiation concluded on day 18. We observed six co-expression clusters of genes during differentiation, and 10uM Pb significantly perturbed two clusters, one involved in cell cycling and DNA repair and the other in protein synthesis. Benchmark concentration analysis identified many genes affected by levels of Pb at or below the current US standard (3.5μg/dL) and these genes were enriched for pathways including stress responses, DNA repair, misfolded protein response, mitosis, and neurotransmitter production. This work highlights potential new mechanisms by which environmentally relevant concentrations of Pb impact gene expression throughout neural differentiation and result in long-lasting implications for neural health and cognition.

## Introduction

Despite major legislative action in the US since the 1970s, environmental lead (Pb) exposure continues to be a persistent public health burden.^1^ Roughly 1.8 million children in the US under the age of 5 have blood lead levels (BLLs) greater than 3.5µg/dL^2^ and roughly half the US population was exposed to levels of Pb above the CDC action level at some point in their childhood.^3^ Pb exposures continue to occur via proximity to industrial sites,^4^ contaminated food^5,6^ and drinking water,^7^ and Pb-containing paint and plumbing from legacy homes.^8^ Exposure to Pb during critical windows of development (e.g., gestation, infancy, and childhood) is associated with adverse neurodevelopmental outcomes, as quantified via learning and memory assessments.^9,10^ This relationship is facilitated by physiological parameters that are specific to these developmental windows, including maternal to fetal transfer of Pb via the placenta,^11^ significant use of hand-to-mouth behaviors during early life,^12^ and an underdeveloped gastrointestinal tract^13^ and blood brain barrier^14^ through childhood.

The mechanisms underlying the neurotoxic effects of Pb have largely been assessed in experimental settings that utilize relatively high exposure levels. Some animal models of Pb’s neurotoxic effects, which have provided evidence of Pb-induced alterations in metabolism and increased oxidative stress^15^, have employed exposure paradigms that resulted in BLLs of up to 30µg/dL,^16^ which is much higher than levels seen at the population level today and for the majority of children exposed in the decades leading up to stricter Pb legislation.^3^ Similarly, much of the *in vitro* work identifying effects of Pb such as DNA damage and mitochondrial dysfunction typically uses exposure levels of 10-25µM (205-518µg/dL).^17^ Many of these studies also used exposure windows that occur outside the developmental process of interest, e.g., before or after cells have differentiated into a neural subtype of interest^18^ as opposed to a continuous exposure throughout differentiation that would more accurately reflect real world exposure scenarios during development. While this foundational work has effectively established high doses of Pb as a hazard in the context of neurodevelopment, characterization of lower, more environmentally relevant and continuous levels of Pb exposure on neurodevelopment is needed.

We seek to investigate the population-relevant Pb exposure effects on the developing nervous system and whether those effects extend beyond mechanisms related to oxidative stress and DNA damage. We previously found that perinatal environmentally relevant Pb exposure led to changes in cell type proportions and altered differentiation in the murine hippocampus.^17^ This work also identified differential gene expression suggesting Pb altered protein folding and stress responses in certain cell types, highlighting the need for additional work in this area. Here we assessed changes in gene expression with exposure to several levels of Pb in the SH-SY5Y model of neural differentiation to address the working hypothesis that exposure to environmentally relevant levels of Pb causes alterations in gene expression and pathways relevant for neurological differentiation, function, and disease, and that effects are differentiation stage-specific.

## Methods

### SH-SY5Y Differentiation and Exposure Conditions

SH-SY5Y cells were purchased from ATCC at passage (P) 28 (Cat. #CRL-2266). Cells were expanded according to manufacturer protocols in a 1:1 mixture of Eagle’s Minimum Essential Media (ATCC, Cat. #30-2003) and F-12 (Thermo, Cat. #11765-054), supplemented with 10% heat-inactivated fetal bovine serum (hiFBS) (Thermo, Cat. #A3840001) and 1% antibiotics (Thermo, Cat. #15140-122). P39 SH-SY5Y cells were differentiated into neuron-like cells using previously published methods.^18^ Briefly, on day (D) 0, 1.2 million cells were seeded onto 15 cm cell culture dishes (Sigma, Cat. #CLS353025) and allowed to adhere overnight. Once attached, cells were maintained for one week in serum-deprivation conditions (2.5% hiFBS) and 10µM retinoic acid (RA) (Sigma, Cat. #R2625). After one week, on D7, cells were split 1:1 onto new 15 cm dishes and maintained for a further three days in 1% hiFBS and 10µM RA. Cells were split once again 1:1 on D10 onto MaxGel extracellular matrix (ECM) (Sigma, Cat. #E0282) in 15 cm dishes and maintained in serum-free Neurobasal media (Thermo, Cat. #21103049) containing brain-derived neurotrophic factor (BDNF) (Thermo, Cat. #10908-010) and dibutyryl cyclic-AMP (db-cAMP) (Fisher, Cat. 16980-89-5). Cells were considered fully differentiated into neuron-like cells on D18. A full description of media conditions can be found in **Table S1**.

SH-SY5Y cells were allowed to begin differentiating in the absence of Pb to reduce the amount of stress on the cells and optimize cell viability. However, once Pb exposure began it was continuous for the remainder of the 18-day protocol. Beginning on D5, differentiating SH-SY5Y cells were exposed to aqueous Pb acetate trihydrate (henceforth referred to as Pb) (Sigma, Cat. #316512) or control conditions. Exposure conditions included 0μM Pb (control), 0.16μM Pb, 1.26μM Pb, and 10μM Pb. These exposure conditions were chosen as they reflect current and historical exposures. 0.16μM Pb corresponds to the blood lead reference value, as of 2024, established by the CDC (3.5μg/dL)^19^ and 1.26μM Pb corresponds to BLLs common in the US prior to the phase out of leaded gasoline and Pb-based paint in the 1970s (26μg/dL).^3^ 10μM Pb corresponds to BLLs > 200μg/dL, which is well above BLLs considered to be tolerable in humans.^20^ Unit conversion for μM to μg/dL was achieved using the molecular weight of Pb (207.2 g/mol) and the equation μM Pb = μg Pb/((0.1dL/L)/207.2 g/mol)). Pb solution was added to cells with each media change (i.e., every 2-3 days). A full illustration of the differentiation and exposure protocol is provided in **Figure 1**.

**Figure 1:**
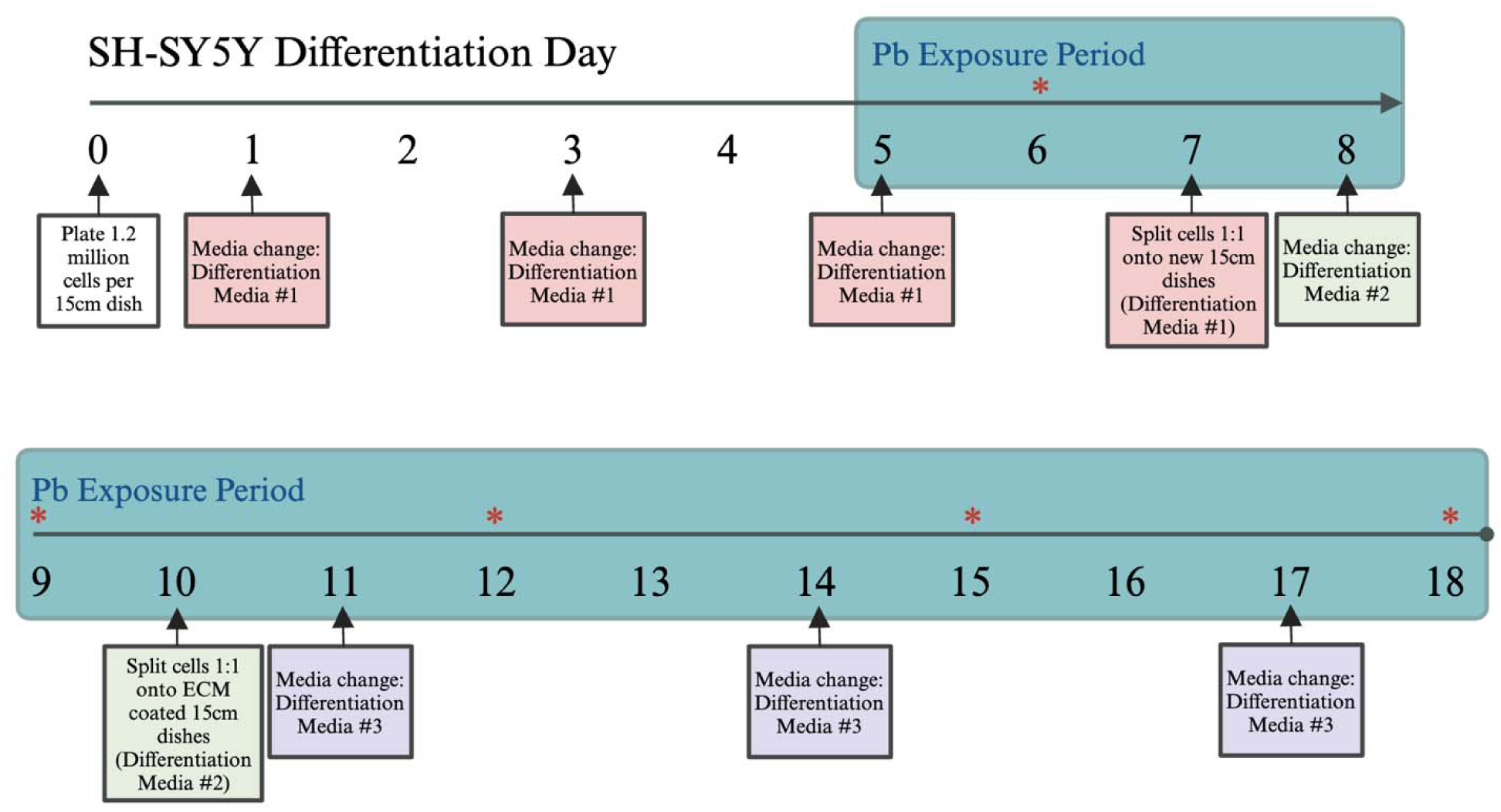
Experimental Workflow. SH-SY5Y cells were differentiated in the presence of 10uM retinoic acid (RA) for 18 days, with lead exposure beginning on Day 5. Cells were collected every three days (indicated by *) to assess changes in gene expression. A complete description of media recipes used can be found in **Table S1**. ECM: Extracellular matrix.

### RNA Extraction and Quantification

Once exposure began on D5, differentiating SH-SY5Y cells were collected every three days (D6, D9, D12, D15, and D18) for RNA extractions. On each day, media was gently removed, and cells were detached from the dish using Tryp-LE Express Enzyme (Thermo, Cat. #1264013) with a 3-minute incubation time at room temperature, or until cells were visibly detached under the microscope. Tryp-LE was quenched with a 5:1 volume of differentiation media and cells were collected via centrifugation at 200g for 5 minutes. Media and Tryp-LE was aspirated and pelleted cells were lysed with 600μL of 1% b-mercaptoethanol (Thermo, Cat. #21985023) diluted in Buffer RLT solution (Qiagen, Cat. #79216), according to manufacturer recommendations. Lysates were stored at -80C until genomic extractions could be completed. A total of = 3 technical replicates were included per exposure condition (n = 4) and time point (n = 5), with a total sample size of N = 60. Lysed samples were homogenized using QIAShredder columns (Qiagen, Cat. #79656) according to manufacturer protocols. RNA was extracted using the AllPrep DNA/RNA/Protein Mini Kit (Qiagen, Cat. #80004) and stored at -80°C until use. RNA concentration was determined using the Invitrogen Quant IT Ribogreen RNA assay kit (Thermo, Cat. #R11490) and standards prepared according to manufacturer recommendations. Standards and samples were plated in black clear-bottom 96 well plate (Corning, Cat. #3603) and were incubated with Ribogreen reagent for 5 minutes, protected from light. Fluorescence was measured on a SpectraMax M5e microplate reader (Molecular Devices, Cat. #89212-400) using the pre-set protocol Ribogreen Assay for Nucleic Assay with SoftMaxPro software version 5.4. 10mg/μL sample dilutions were prepared for each sample based on quantification results.

### plexWell cDNA Preparation and Quality Control Assessment

RNA was sequenced via the plexWell (SeqWell) plate-based method, adapting our previously developed protocol.^21^ For cDNA preparation from RNA, briefly, 1μL of 10ng/μL RNA was added to a 96 well plate (Dot Scientific, Cat. #951-PCR-B) for oligo annealing at 72°C for 10 minutes. cDNA was then amplified at 50°C for 30 minutes, 98°C for 3 minutes, followed by 12 cycles of 98°C for 20 seconds, 67°C for 20 seconds, and 72°C for 6 minutes, in the C1000 Thermal Cycler (Biorad, Cat. #1851196) and samples were held at 4°C until purification. cDNA was purified using an equivalent amount of MAGwise Paramagnetic beads (SeqWell, Cat. #MG10000) and a binding time of 5 minutes. Samples were placed on a 96 well plate magnet, supernatant was removed, and beads were washed once with 80% ethanol (Spectrum, Cat. #E10258). cDNA was eluted in 20μL of 10mM Tris (Thermo, Cat. #J22638-AE) and stored at - 20°C until quantification.

cDNA concentrations were determined using the Quant-iT Picogreen dsDNA assay kit (Thermo, Cat. #P7589) according to manufacturer protocols. Samples and standards were plated in a 96 well plate and Picogreen solution was added at a volume of 100μL/well, followed by an incubation at room temperature protected from light. Fluorescence was measured on the SpectraMax M5e using the pre-set protocol Picogreen assay for Nucleic Assay with SoftMaxPro software version 5.4. A plexWell-provided global dilution factor calculator was used to determine the final volume of cDNA to ensure a minimum input of 5ng per sample.

### plexWell Library Preparation and RNA Sequencing

6µL of cDNA per sample at approximately 1.7ng/μL was used for library preparation. cDNA was added to a hard skirted sample barcode plate provided in the plexWell LP384 Library Preparation Kit (SeqWell, Cat. #LP384X). Each sample was labeled with an i7 index Sample Barcode via PCR conditions of 55°C for 15 minutes, followed by 68°C for 10 minutes. After i7 tagging, 18μL of each barcoded samples were pooled, and an equivalent volume of MAGwise paramagnetic beads were added and cDNA was allowed to bind for 5 minutes. Sample pools were placed on a magnetic stand and beads allowed to pellet, followed by two washes with 80% ethanol. Barcoded cDNA was eluted in 40μL of 10mM Tris. Post pooling, quantification was achieved using the Picogreen assay described above to ensure the eluate was within the protocol-recommended range (3.6-6ng/μL).

The 2 sample barcoded pools were then labeled with an i5 index (i.e., Pool Barcode), using the tagmentation reaction (55°C for 15 minutes, 68°C for 10 minutes). i5-tagged pools were cleaned using the protocol described above and each pool was eluted in 24μL of 10mM Tris. Purified pools were amplified at 72°C for 10 minutes, 95°C for 3 minutes, 8 cycles of 98°C for 30 seconds, 64°C for 15 seconds, and 72°C for 30 seconds, followed by 72°C for 3 minutes, after which samples were held at 4°C until use. Following amplification, library products were diluted using 205μL of 10mM Tris. 200μL of the diluted product was purified using 0.8 equivalents of MAGwise paramagnetic beads, a 5-minute binding time, pelleting on a magnetic stand, and two washes with 80% ethanol. Purified multiplexed library was eluted with 30μL of 10mM Tris and stored short term at -20°C. Library QC was performed on the Agilent Bioanalyzer (High Sensitivity DNA 5000 kit) at the UM Advanced Genomics Core.

Libraries were sequenced on the Illumina NovaSeq 6000 at the UM Advanced Genomics Core, using 151-bp paired end reads to an average depth of great than 30 million reads per sample. FastQC reads were demultiplexed back into individual samples based on the i7 and i5 indices. Sequencing data was transferred to the UM Great Lakes high-performance computing cluster for analysis. Sequencing read quality was assessed via *FastQC*^22^ and *MultiQC*^23^. Reads were aligned to a splice junction-aware build of the human genome (GRCh38) using *STAR*.^24^ Gene counts per million were computed using *featureCounts*,^25^ where multimapping and multi-overlapping reads were not counted.

### Differential Gene Expression Analysis

Read count matrices were loaded into *edgeR*^26^ in R (v.4.3.3). Genes with low expression were excluded from downstream analysis using the default settings of *filterByExpr*. Library sizes were normalized using *calcNormFactors* and dispersion estimated using *estimateDisp*. Differential gene expression between each Pb condition (n = 3 replicates) and relevant controls (n = 3 replicates) at each time point was calculated using quasi-likelihood negative binomial generalized log-linear modeling. Genes were considered differentially expressed between a Pb exposure condition and a corresponding control, or between time points, with a |fold-change in expression| > 2 (|log_2_FC| > 1) and a Benjamini–Hochberg^27^ false discovery rate (FDR)-adjusted p-value of < 0.05. We conducted an additional analysis in the control cells, evaluating differential gene expression as differentiation progressed in the absence of Pb, wherein D6 (the earliest time point captured) was used as the reference. A |fold-change in expression| > 2 (|log_2_FC| > 1) and an FDR-adjusted p-value of < 0.05 were used. We visualized differential gene expression using volcano plots.

### Cluster Analysis

Read counts were imported into *clust*, a Python-based method of identifying co-expressed gene clusters from gene expression data based on biological expectations of gene co-expression,^28^ using v.1.18.0. *Clust* generates gene clusters with lower levels of dispersion than other methods of co-expression analysis and is more reliably reproducible, such as partitioning or partial clustering. We visualized expression patterns within a cluster using spaghetti plots of differentiation day versus gene expression, with a line for each gene. Gene cluster profiles and cluster objects containing lists of genes represented in each cluster were imported into R for integrated analysis with differential gene expression results. Using a hypergeometric test, we assessed whether any clusters identified in this analysis were enriched for Pb-responsive genes (Pb-responsive genes defined as differentially expressed at any time point in the 10μM Pb condition, relative to control cells, with a |logFC| > 1 and FDR < 0.05). Gene set enrichment testing was conducted for each cluster using Go.db and “org.Hs.eg.db” from Bioconductor.

### Benchmark Concentration Modeling and Gene Set Enrichment Analyses

Gene-level benchmark concentration (BMC) analysis was conducted using BMDExpress v.3.2 according to best practices outlined in online documentation.^29,30^ This software was developed by the National Institute of Environmental Health Sciences (NIEHS). Normalized counts per million reads were imported and prefiltered using one-way analysis of variance (ANOVA). This prefilter step identified genes with significant increasing or decreasing expression with Pb concentration with significance threshold set at p < 0.05. Filtered data was modeled in BMDExpress with Hill, power, linear, polynomial, and exponential models, and the best fit model was chosen by the software for each concentration-response relationship per gene using nested chi square. Hill models were flagged if the “k” parameter was smaller than 1/3 of the lowest positive concentration but not omitted from the data. The best benchmark concentration represents the concentration of Pb that produces a significant change in gene expression. We calculated the median best benchmark concentration and the minimum best benchmark concentration across all genes for each time point in this experiment.

We conducted enrichment analyses as well as “Defined Category” analyses to assess the dose response relationship to the Gene Ontology (GO)^31^ and the Molecular Signatures (MSigDB) “hallmark” gene sets.^32^ We supplemented the hallmark gene set with additional gene sets from MSigDB, including Lein_Neuron_Markers,^33^ Lee_Neural_Crest_Stem_Cell,^34^ and Le_Neuronal_Differentiation.^35^ Benchmark doses with values above the highest tested concentration (10µM) were excluded from individual gene and functional classification analyses. Pathway analyses results were deemed significant with a Fisher’s exact two-tailed p-value of <0.05 and results were visualized using the R package *ggplot*. All original expression data, ANOVA filtered results, BMC results, and BMDExpress3 analysis is included in a .bm2 file available in supplementary information.

## Results

### Differential Gene Expression with Lead Exposure

Differential gene expression was assessed in differentiating SH-SY5Y cells exposed to environmentally relevant doses of Pb, delivered via cell culture media. For each time point, control (0μM Pb) cells were used as the reference group against which the low (0.16μM Pb), medium (1.26μM Pb), and high (10μM Pb) conditions were compared. Results are illustrated via volcano plots, where genes labeled in red were those with significant changes in expression in exposed cells relative to control (**Figure 2**) and are summarized in **Table 1**. A full summary of all differential expression results is provided in **Table S2**. D6 of differentiation corresponds to 24 hours after the onset of Pb exposure (D5) and no significant changes in gene expression were detected at any dose (FDR > 0.05) (**Figure S1A-C**). The second time point assessed, D9, corresponded to four days of continuous Pb exposure as well as 48 hours after the first split of cells onto new plates. On D9, we observed minimal changes in gene expression at the low and medium Pb exposure conditions. 0.16μM Pb exposure resulted in a significant increase in the expression of *HMOX1* (logFC = 2.44, FDR = 0.002) (**Figure 2A**). Similar results were observed with 1.26μM Pb exposure (*HMOX1*, logFC = 2.22, FDR = 2.29×10-3), along with additional significant increase in the expression of *NQO1* (logFC = 1.13, FDR = 4.86×10-5) and *OSGIN1* (logFC = 1.51, FDR = 3.91×10-2), and significant decreases in the expression of *KRT18* (logFC = -1.39, FDR = 4.45×10-5) and *COL3A1* (logFC = -1.3, FDR = 5.16×10-5) (**Figure 2B**). 10μM Pb exposure resulted in a distinct but limited set of upregulated genes, including *GDF15*, *KLHDC8A*, and *LCIIAR* and 70 significantly downregulated genes, including *KRT18* and *COL3A1* (**Figure 2C**).

**Figure 2:**
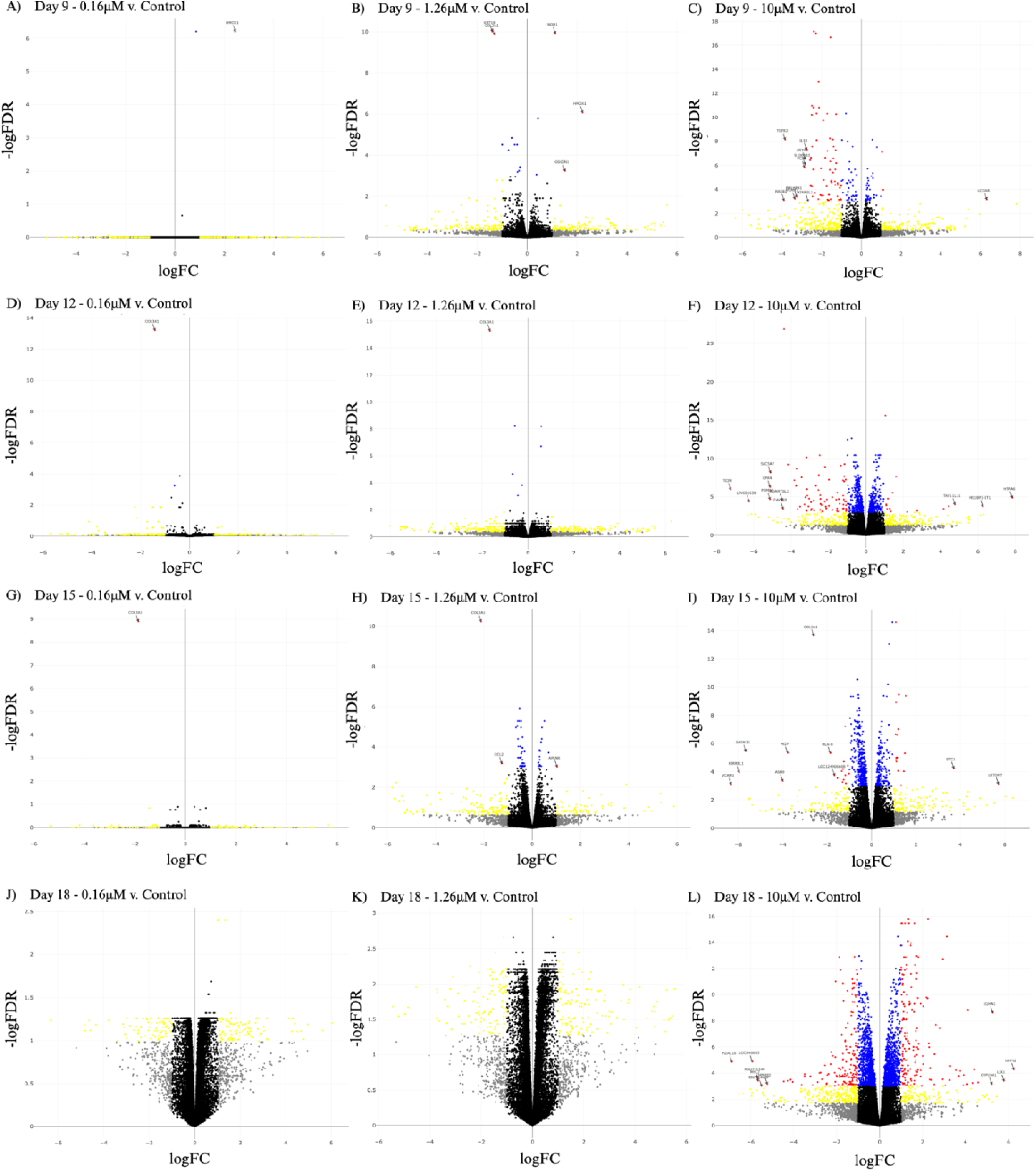
Differential Gene Expression Associated with Lead Exposure During SH-SY5Y Differentiation. Differential gene expression during differentiation was calculated for each dose (0.16_μ_M, 1.26_μ_M, and 10_μ_M Pb) relative to control cells at each time point (**Day 9: A-C**, **Day 12: D-F**, **Day 15: G-I**, **Day 18: J-L**). The x-axis shows the log_2_ fold change differential gene expression between lead treatment and control. Higher x-axis values represent higher expression with lead treatment. The y-axis shows the -log_10_ false discovery rate adjusted p-values. Higher y-axis values represent more significant associations. Significantly differentially expressed genes (|log_2_FC| > 1, FDR < 0.05, p-value < 0.05) denoted in red, differentially expressed genes (|log_2_FC| > 1, p-value < 0.05) and significant genes (FDR < 0.05 and p-value < 0.05) denoted in yellow and blue, respectively. Genes with insignificant changes in expression (|log2FC| > 1, p-value and FDR > 0.05) denoted in grey and genes with no detectable change in expression between conditions (|log_2_FC| < 1, p-value and FDR > 0.05) denoted in black. Corresponding counts of differentially expressed genes at each time point can be found in **Table S2**. logFC: log_2_ fold change differential expression between Pb treatment and control. FDR: false discovery rate adjusted p-values.

**Table 1:** Summary of Differentially Expressed Genes Associated with Lead Exposure During SH-SY5Y Differentiation. Total counts of the number of significantly (|logFC| > 1, FDR < 0.05) differentially expressed genes at each time point provided, along with the total number of genes that displayed increased expression (Up) and decreased expression (Down) with lead exposure. logFC: log_2_ fold change differential expression between Pb treatment and control. FDR: false discovery rate adjusted p-values

| | 0.16$\mu\text{M}$ v. Control | | | 1.26$\mu\text{M}$ v. Control | | | 10$\mu\text{M}$ v. Control | | |
| --- | --- | --- | --- | --- | --- | --- | --- | --- | --- |
|  | Up | Down | Total | Up | Down | Total | Up | Down | Total |
| **Day 6** | 0 | 0 | 0 | 0 | 0 | 0 | 0 | 0 | 0 |
| **Day 9** | 1 | 0 | 1 | 3 | 2 | 5 | 3 | 70 | 73 |
| **Day 12** | 0 | 1 | 1 | 0 | 1 | 1 | 17 | 71 | 88 |
| **Day 15** | 0 | 1 | 1 | 1 | 2 | 3 | 17 | 19 | 36 |
| **Day 18** | 0 | 0 | 0 | 0 | 0 | 0 | 171 | 160 | 331 |

The D12 time point corresponded to 7 full days of Pb exposure and 24 hours after the addition of terminal differentiation factors, including BDNF. As with D9, in D12 a limited set of genes were differentially expressed in the 0.16μM and 1.26μM Pb conditions, as the only significant change in gene expression was the downregulation of *COL3A1* (**Figure2D-E**). 10μM Pb exposure led to the significant upregulation of 17 genes and the downregulation of 71 genes. This latter set again included *KRT18* (logFC = -1.14, FDR = 9.52×10-5) and *COL3A1* (logFC = -4.37, FDR = 2.1×10-12) (**Figure 2F**). This pattern was replicated on D15, with the significant downregulation of *COL3A1* expression under the 0.16μM and 1.26μM Pb conditions, however the 1.26μM Pb condition also resulted in the significant downregulation of *CCL2* (logFC = -1.26, FDR = 0.0041) and upregulation of *APLNR* (logFC = 1.02, FDR = 0.04) (**Figure 2G-H**). On D15, 10μM Pb exposure led to the significant down and upregulation of 17 and 19 genes, respectively (**Figure 2I**). As differentiation concluded on D18, significant changes in gene expression were nonexistent for the 0.16μM and 1.26μM Pb conditions (**Figure 2J-K**). 10μM Pb exposure caused the greatest amount of differential gene expression of any exposure-time point combination, with 160 genes significantly downregulated in their expression and 171 significantly upregulated (**Figure 2L**).

### Differentiation Alters Patterns in Gene Expression Among Unexposed Cells

To help contextualize the effects of Pb throughout differentiation, we first characterized the gene expression changes which occurred in control cells across the time course. These analyses identified significant changes that were differentiation stage-dependent, when using D6 as the reference group as it was the first time point collected in these experiments (**Figure S2**, **Table S3**). There was an increase in significantly expressed genes as differentiation progressed with 98 on D9 relative to D6 (63 upregulated/35 downregulated), 2160 on D12 (1437/723), 2598 on D15 (1743/864), and 3989 on D18 (2377/1612) (**Table S4**). We performed cluster analysis using *clust* to ascertain whether groups of co-expressed genes displayed similar patterns of expression over the course of SH-SY5Y differentiation and by Pb exposure. Six gene clusters were identified (C0-C5, **Figure 3A**) and pathway analysis of gene members of each cluster via the Gene Ontology (GO) database revealed significant enrichment for five of the six clusters (**Figure 4**).

**Figure 3:**
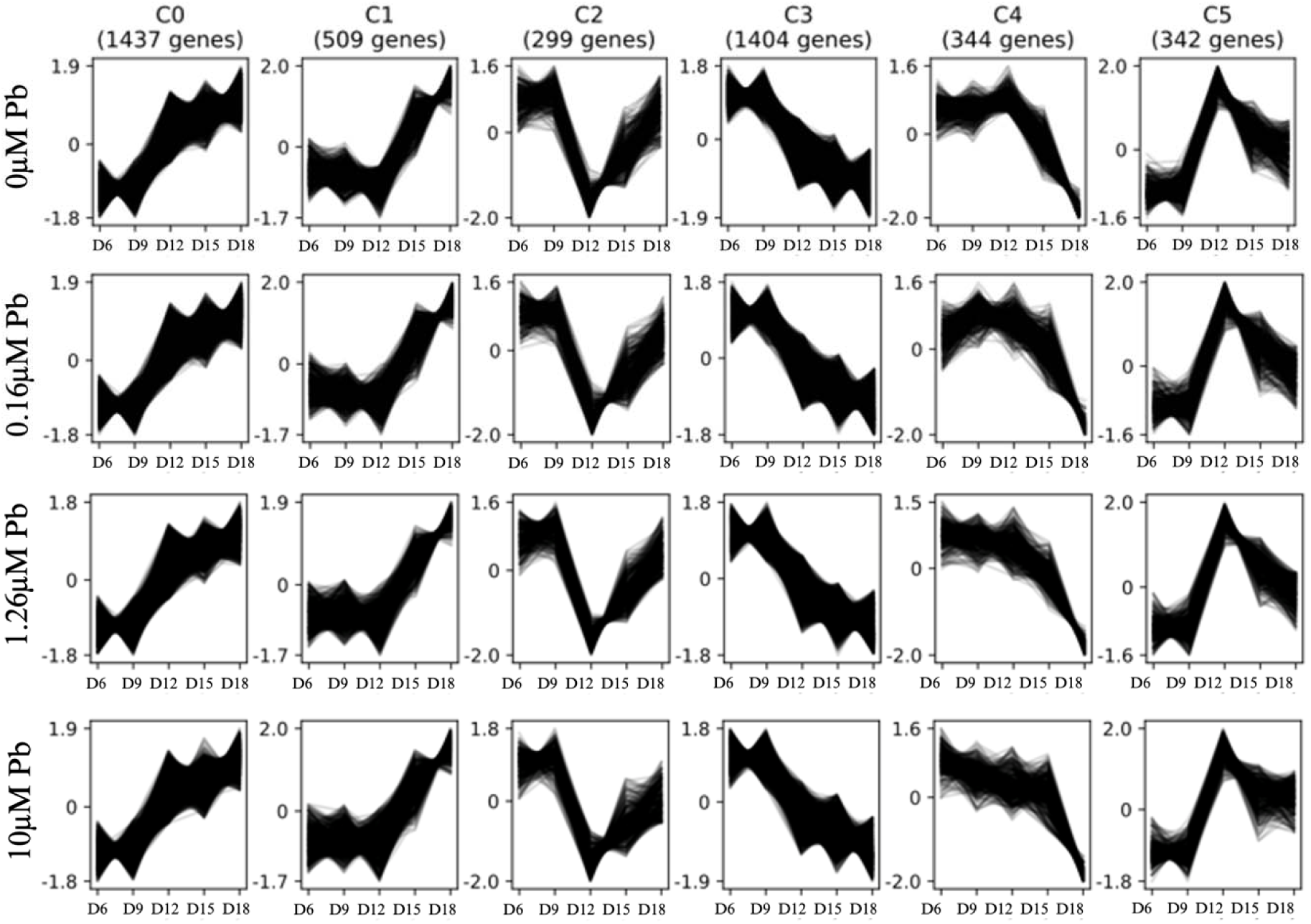
Cluster Analysis of Gene Count Matrices in Control and Exposd SH-SY5Y Cells During Differentiation. Six gene clusters identified (C0-C5) over the course of differentiation (Days 6 – 18) with the number of genes identified for a given cluster in parentheses. The x-axis shows differentiation day. The y-axis represents normalized gene expression values for each gene. Each line represents expression over time for a given gene.

**Figure 4:**
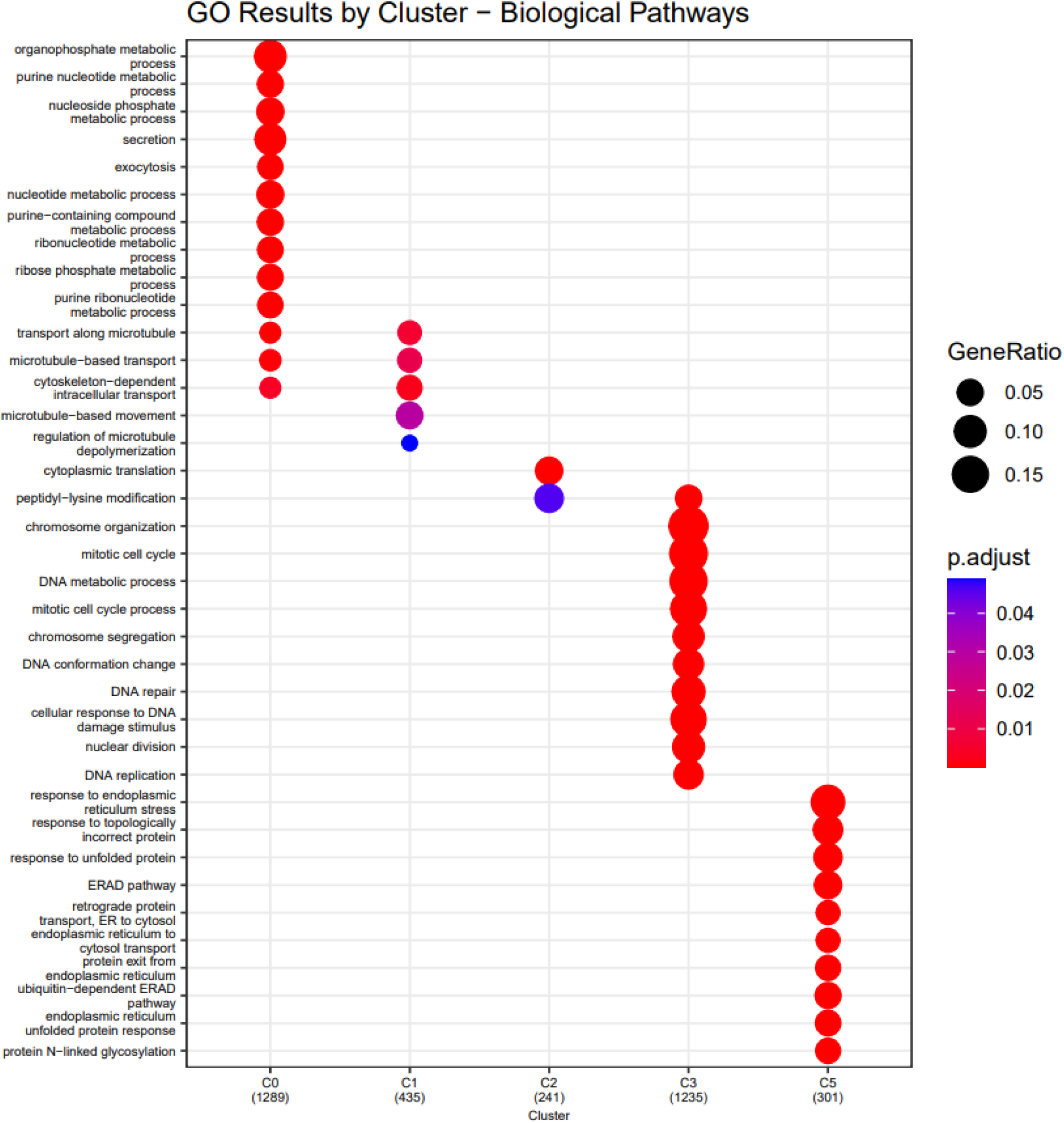
Gene Ontology (GO) Enrichment of Gene Clusters. Lists of genes in each cluster identified in Figure 3 were subjected to pathway analysis. The x-axis shows each cluster (C0 – C5) and the number of genes annotated to that cluster. The y-axis shows the biologic pathway that was enriched. The size of the circles corresponds to the gene ratio (defined as the number of differentially expressed genes divided by the number of genes associated with that GO term). The color of the circle corresponds to the false discovery rate adjusted p value, with more significant values bluer on the color spectrum and less significant values shown in red.

Cluster 0 (C0) was enriched for genes involved in cellular metabolism and expression of this set steadily climbed throughout differentiation. C1, enriched for genes related to cellular structure, was somewhat stagnant during early differentiation but increased rapidly starting on D12 and through the remainder of the differentiation process. C2 had modest enrichment of genes involved in translation, and this cluster had relatively high expression during early differentiation, followed by a steep decline between D9 and D12, after which expression began to climb again through D18. Genes in C3 were primarily relevant to cellular replication and DNA regulation, and this cluster had elevated expression early in differentiation followed by a consistent decline through D18. Finally, genes in C5 were enriched for those involved in protein regulation, endoplasmic reticulum stress, and the unfolded protein response, and their expression gradually increased between D6 and D12, after which they declined again as differentiation concluded (**Figure 3A and 4**).

### Differentiation Genes Overlap with Pb-Response Genes

Two clusters were significantly enriched with Pb-responsive genes. C3, with a total of 1404 genes in the cluster, contained 55 genes that were significantly affected by 10μM Pb exposure (p = 0.0032), and C5, with 342 genes in the cluster, contained 30 genes that were significantly impacted by 10μM Pb exposure (p = 1.53×10-8) (**Table 2**). Given these results, we refined our differential expression analysis with a focus on the expression patterns of 10μM Pb-responsive genes present in C3 and C5. We found a consistent pattern across time points, in that Pb-responsive genes overlapping with C3 were nearly all significantly downregulated with Pb exposure while those overlapping with C5 were much more likely to be upregulated (**Figure S3**).

**Table 2:** Test of Enrichment of Differentially Expressed Genes in Identified Gene Clusters. We tested genes in each of the six gene clusters for enrichment in the Pb gene signature list (those genes that displayed differential expression at any dose and on any day) using a hypergeometric test (*phyper*). Results are presented by Cluster and include the number of genes in a given cluster, the number of significant DEGs (|log_2_FC| > 1, p < 0.05) that overlapped with that cluster, and unadjusted p-value calculated using # of Pb signature genes = 472 for all comparisons. Significant results are highlighted in red. DEGs: Differentially expressed genes. Log_2_FC: log_2_ fold change comparing gene expression with lead treatment to control.

| Cluster | # of Genes in Cluster | # Overlapped with Pb Signature | p-value |
| --- | --- | --- | --- |
| 0 | 1437 | 30 | 0.94 |
| 1 | 509 | 6 | 0.99 |
| 2 | 299 | 9 | 0.41 |
| 3 | 1404 | 55 | 0.003 |
| 4 | 344 | 2 | 0.99 |
| 5 | 342 | 30 | $1.53 \times 10^{-8}$ |

### Benchmark Concentration and Gene Set Enrichment Analysis

We performed benchmark concentration analysis to characterize the concentration-response relationship of Pb exposure on gene expression at each time point using BMDExpress3. Benchmark concentrations were estimated from the Pb dose-response model at each time point for each gene. The total number of genes impacted by Pb across the concentration range increased with duration of exposure, with Pb inducing concentration-response differences at or under 10μM Pb exposure in over 4,500 genes by Day 18 (**Figure 5**). Concentration-response fit was assessed via examination of the best fit Pb concentration-response curves for individual genes at each time point. The entirety of benchmark concentration response data for each differentially expressed gene across all time points is included in the supplementary .bm2 file and an overview of best benchmark concentrations estimated for each gene across differentiation is illustrated in **Figure 6** and **Table 3**, with the median best benchmark concentrations falling between 2.92μM and 3.34μM.

**Figure 5:**
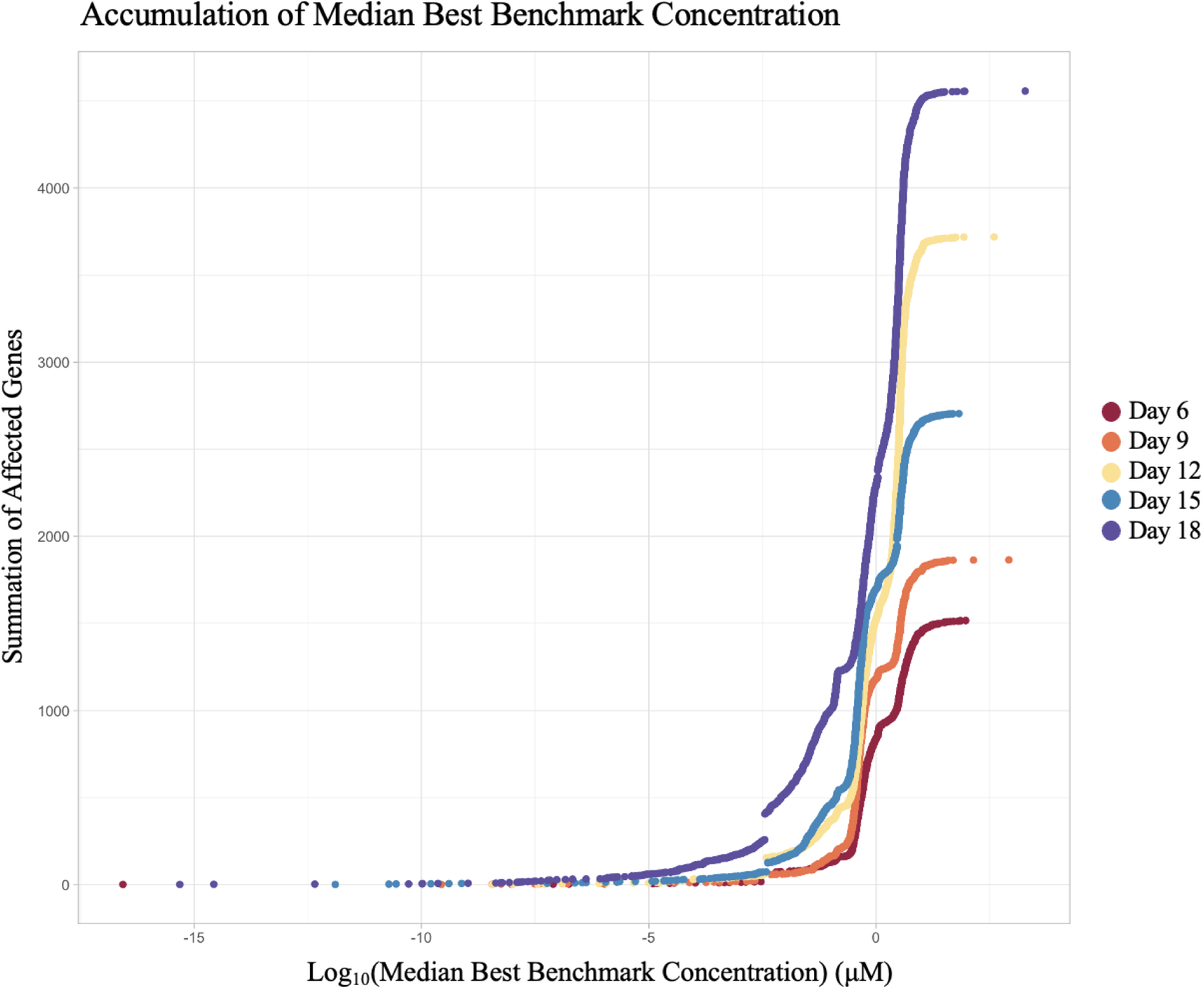
Accumulation Plot of Differentially Expressed Genes with Increasing Lead Dose During SH-SY5Y Differentiation. The x-axis shows the log_10_-transformed median best benchmark concentration. The y-axis shows the number of genes affected by a given dose. The colors of the points correspond to the day of differentiation (Day 6, 9, 12, 15, and 18).

**Figure 6:**
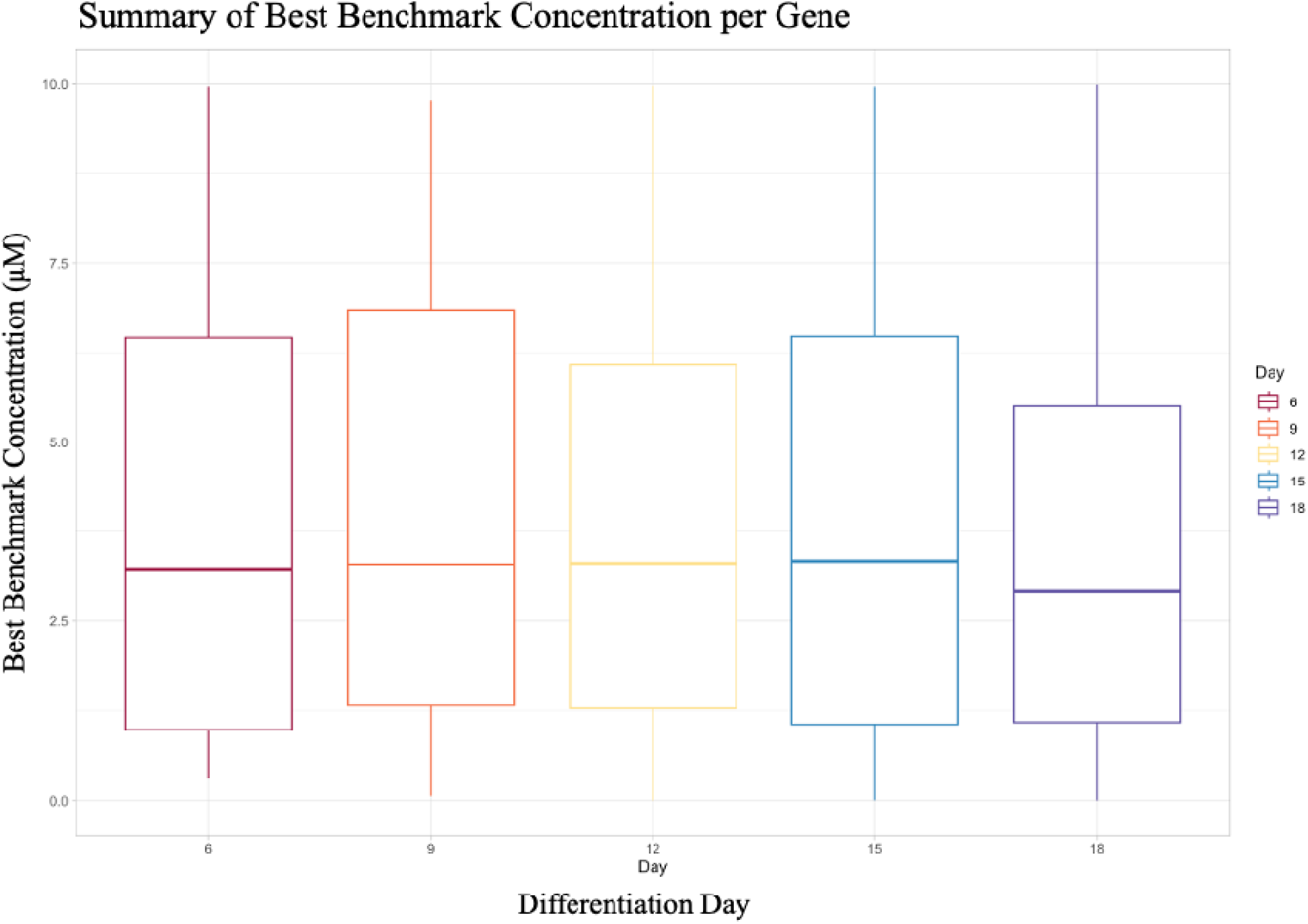
Summary of Best Benchmark Concentration of Lead During SH-SY5Y Differentiation. Best benchmark concentrations of lead (Pb) exposure for each gene are summarized by timepoint in boxplots, wherein the limits of each box represent the interquartile range, the midline represents the median best benchmark concentration, and lower and higher tails extend to the minimum and maximum best benchmark concentrations, respectively. For the purposes of this analysis, best benchmark concentrations greater than 10_μ_M Pb were not considered.

**Table 3:**
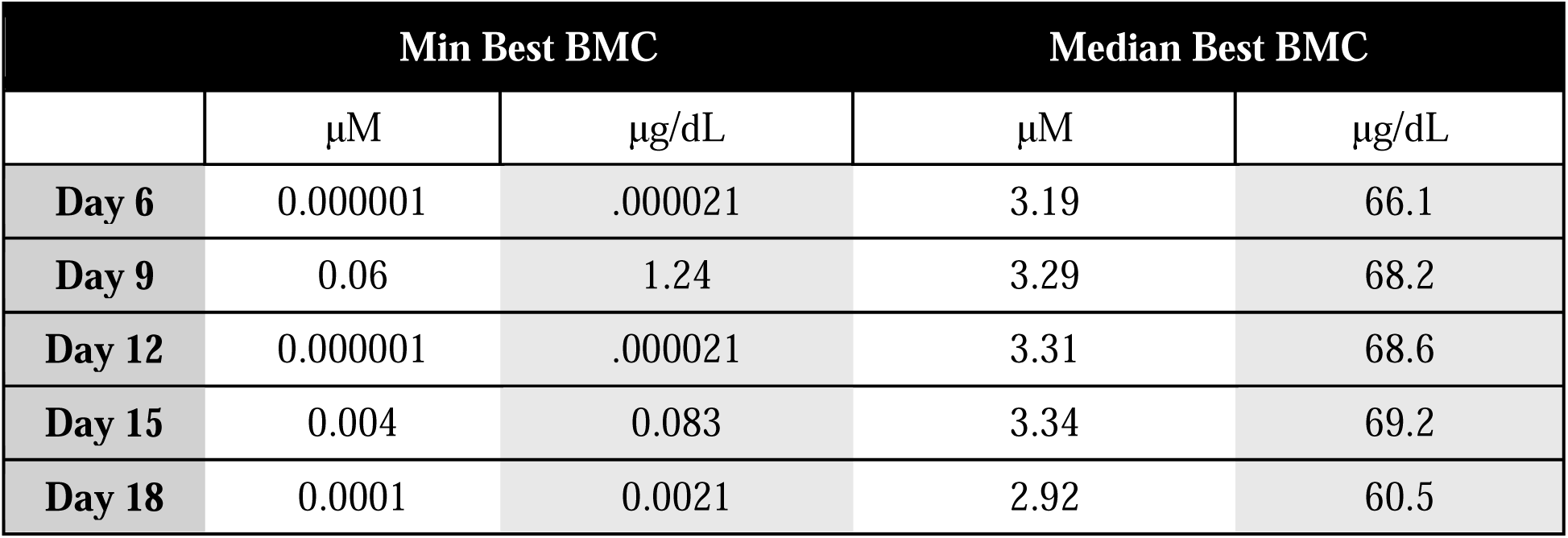
Summary of Minimum and Median Best Benchmark Concentrations of Lead During SH-SY5Y Differentiation. BMD: benchmark concentration.

Following best benchmark concentration analysis, we used BMDExpress3 to conduct gene set enrichment testing using the built-in Functional Classification function and two background datasets. A complete summary of functional classification data is included in the supplementary .bm2 file. The number of significantly enriched gene sets from the GO Biological Pathways database increased as SH-SY5Y differentiation progressed (**Figure 7A, Table S4**. On Day 9, Pb-responsive genes were enriched for those involved in deoxyribonucleoside biosynthetic processes and neural processes including cranial ganglion and neural projection development. As a whole, we observed a decrease in the expression of these genes, all at environmentally relevant levels of Pb exposure (0.456-0.462μM Pb). By Day 12 we observed a shift in enrichment patterns, with Pb-responsive genes associated with pathways related to cell division, cell growth, and metabolism. Pb-responsive genes involved in these pathways were largely downregulated and affected at relatively high levels of exposure (1.63-3.2μM Pb). Notable pathways significantly associated with Pb-responsive genes on Day 15 included deoxyribonucleoside biosynthesis and DNA replication, at concentrations ranging from .04-1.29μM Pb and a continuation of downregulated expression seen at earlier time points. A notable reversal of this pattern was that of cholesterol biosynthesis, as Pb-responsive genes involved in this pathway were upregulated in their expression. As differentiation concluded on Day 18, Pb-responsive genes were enriched for those associated with DNA replication and repair and cellular replication (largely downregulated), and neurotransmitter synthesis (upregulated) with the vast majority affected at environmentally relevant concentrations (0.03-1.58μM Pb).

**Figure 7:**
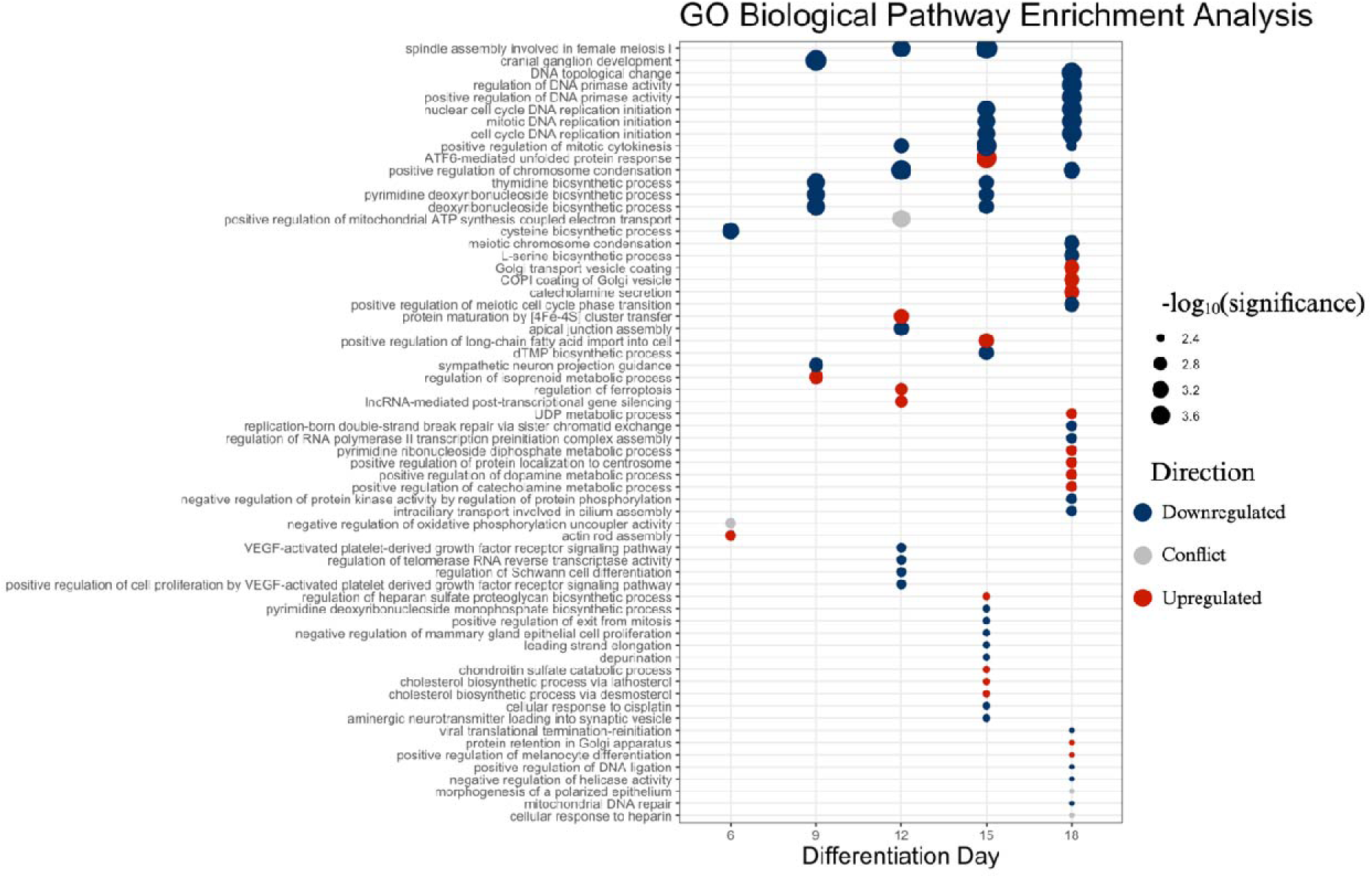
Gene Ontology Biological Pathway Analysis of Differentially Expressed Genes. Gene expression data was imported into BMDExpress and filtered for significantly differentially expressed genes (|logFC| > 1, FDR < 0.05) using One-Way ANOVA pre-filter followed by Functional Classification and the Gene Ontology function. Pathways significantly enriched for Pb-responsive genes (p < 0.05), with the size of each dot proportional to significance and color indicating whether the majority of Pb-affected genes in a given pathway were downregulated (blue) or upregulated (red). If there was no majority, the relationship was classified as conflicting (grey). Highly significant results (p<0.05, Pb-responsive genes:total number of genes in pathway ratio > 0.7) are included here with all results included in the supplemental .bm2 file.

A second enrichment analysis was performed within BMDExpress 3, beginning with the “hallmarks” gene set from MSigDB, supplemented with gene sets related to neural stem cells and markers (**Figure 8A-B**). Using the supplemental gene sets, we found neuro-related markers to be most affected early on in differentiation (Day 6), with a median BMC of 0.55μM or 11.39μg/dL Pb, alongside upregulation of hallmarks of stress responses (e.g., UV and reactive oxygen species, median BMC = 0.68-2.81μM Pb) and downregulation of genes involved in cell cycling (e.g., E2F targets and G2M checkpoint, median BMC = 0.35-0.37μM Pb). By Day 12, we observed decreased expression of genes typically upregulated during neural differentiation at a median BMC of 0.64μM Pb. We also observed a downregulation of genes linked to stress responses (e.g., gamma, interferon, and inflammatory responses) and a corresponding increase in genes associated with DNA repair, the p53 pathway, and misfolded protein responses, all at median BMCs of 0.2-2.33μM Pb. Day 15 and 18 continued the trend of downregulation of genes involved in cell cycling (median BMC = 0.13-0.42μM Pb), alongside increased expression of genes involved in apoptosis and unfolded protein response on Day 15 at moderate Pb levels (median BMC = 0.4-0.67μM Pb) and downregulation of those related to cell cycling and DNA repair on Day 18 (median BMC = 0.049-1.38μM Pb).

**Figure 8:**
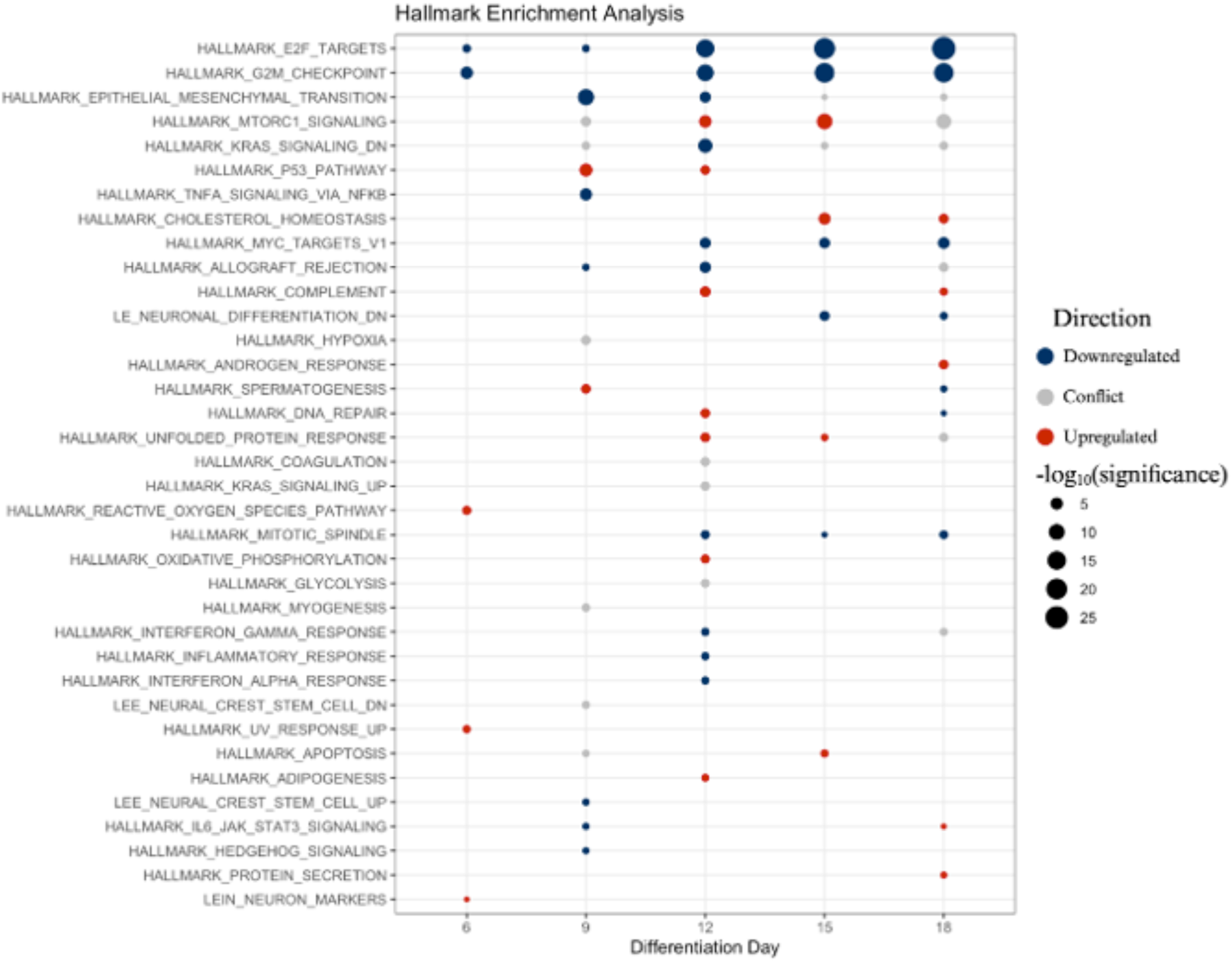
Molecular Signatures Database and Supplemental Dataset Analysis of Differentially Expressed Genes. Gene expression data was imported into BMD Express and filtered for significantly differentially expressed genes (llogFCI> 1, FDR< 0.05) using One-Way ANOVA pre-filter followed by Functional Classification and imported Hallmarks and supplemental datasets from Molecular Signatures Database (MSigDB). Pathways significantly enriched for Pb-responsive genes (p < 0.05), with the size of each dot proportional to significance and color indicating whether the majority of Pb-affected genes in a given pathway were downregulated (blue) or upregulated (red). If there was no majority, the relationship was classified as conflicting (grey). Highly significant results (p<0.05, Pb-responsive <u>genes:total</u> ration> 0.6) are included here with all results included in the supplemental .bm2 file.

## Discussion

### Differential Gene Expression in Response to Lead is Differentiation Stage-Specific

Here we demonstrate time course-specific effects of Pb exposure on gene expression during SH-SY5Y differentiation into neuron-like cells. While most observed changes occurred in the highest concentration (10μM Pb), there were notable and consistent changes in low (0.16μM Pb) and medium (1.26μM Pb) conditions as well (**Figure 2**). Expression of *COL3A1* was decreased by Pb exposure across concentrations beginning on D9. *COL3A1* encodes the alpha-1 chain of type III collagen, with relevance to connective tissue,^36^ wound healing,^37^ and cortical lamination and neuronal migration.^38^ *COL3A1* loss leads to cortical delamination in mice^39^ and *in vitro* work revealed abundant presence of this transcript and its product in differentiated neurites.^40^ Given the suppression of *COL3A1* expression with Pb exposure, future work should examine whether this relationship drives a reduction in neural migration.^41^

An advantage our approach was quantifying gene expression at multiple points during differentiation with continuous Pb exposure. Much of the work assessing Pb effects on neural differentiation *in vitro* use paradigms where exposure is limited to before^42^ or after^43^ differentiation occurs. Results from these studies are challenging to extrapolate to effects in human populations as the majority of Pb exposures are chronic, from the transfer of maternal stores during gestation^44^ or contaminated environments.^45^ By profiling across time and concentrations, we observed gene expression changes in the middle of SH-SY5Y differentiation (downregulation of *COL3A1* on D9, D12, and D15, and *KRT18* on D9; upregulation of *HMOX1* and *NQO1* on D9 and *APLNR* on D15) with low and medium Pb exposures, all of which disappeared as differentiation concluded on D18 (**Figure 2**). Lack of *KRT18* in the hippocampus is linked with apoptosis and ferroptosis,^46^ cellular processes linked with memory impairment.^47^ Overexpression of *HMOX1* and *NQO1* suggests a response to oxidative stress,^48^ aligning with an abundance of work linking Pb exposure to this outcome in the nervous system.^49,50^ While changes in expression of these genes were transient, they may cause longer-lasting changes in cellular function. Future investigations which focus on these effects may elucidate novel mechanistic insights.

### Lead Exposure Impacted Specific Clusters of Genes in a Cohesive Manner

Cluster analysis of co-regulated genes during differentiation revealed six distinct groups, with two of the six (C3 and C5) enriched for Pb-responsive genes. Pb-sensitive C3 was enriched for genes involved in cellular division and DNA repair (**Figure 4**). In human populations, Pb exposure has been associated decreased expression of genes related to DNA repair,^51,52^ and *in vitro* evidence suggests there may be interactions between Pb and epigenetic machinery that prevent DNA repair mechanisms from operating properly.^53^ Similarly, we observed a trend of decreased expression in C3 genes that were also Pb-responsive, suggesting Pb suppresses DNA repair response. This relationship is also interesting as the depressive effect of Pb on genes in the C3 cluster was more pronounced as differentiation progressed towards D18 (**Figure S3**), and may be due to several mechanisms, such as the accumulation of Pb in the cells^54^ and corresponding increases in oxidative stress, a fatigue in the DNA repair system over time,^49^ or a combination of these or other variables that change over time.

C5 was enriched for Pb-responsive genes and pathway analysis revealed this cluster was related to protein production and endoplasmic reticulum function (**Figure 4**). Generally, Pb-responsive genes in C5 demonstrated increased expression with exposure. These results align with work demonstrating relationships between protein misfolding and oxidative stress^55^ and DNA damage^56^, both linked to Pb.^49,57^ Prolonged endoplasmic reticulum stress and protein misfolding, and resulting aggregates, have been heavily implicated as hallmarks of neurodegenerative disease, such as Alzheimer’s disease.^58^ Exposure to toxicants such as Pb, that put stress on this cellular function specifically, increase susceptibility outcomes related to neurodegeneration due to protein misfolding events and prolonged stress on the endoplasmic reticulum.^59^

### Current and Historical Concentrations of Lead Affect Gene Expression

We utilized BMDExpress to determine benchmark concentrations of Pb exposure on individual genes across timepoints (**Figure 5**). The median best benchmark concentration (calculated across all genes with a best benchmark concentration lower than the highest dose tested in this study, i.e., 10μM Pb) for all time points included in this study was around 3μM (61μg/dL). While this concentration is higher than exposure levels typically seen today in the US (but not unheard of worldwide^60^) BLLs greater than 25μg/dL are still reported for workplace exposures^61^ as well as in children living in environments with high Pb contamination.^62^

BLLs greater than 60μg/dL were much more common in the US prior to the phase-out of Pb-based paint and gasoline.^3^ In fact, a BLL of 60μg/dL was the threshold used by the CDC prior to 1960 to indicate intervention was necessary.^63^ Thus, we hypothesize that individuals who were children prior to 1960 in the US were much more likely to have BLLs high enough to significantly impact gene expression, which may correspond to alterations in neurodevelopment.^64^ Additionally, in light of the work on the effects of Pb on protein misfolding, these results add to the discussion that developmental exposure to Pb may contribute to neurodegenerative disease^65^ and could partially explain increases in neurodegenerative disease rates.^66^ Individuals who were children in the mid-20^th^ century are entering into their 7^th^ and 8^th^ decades today, phases of life during which neurodegenerative disease increases in severity and often is diagnosed for the first time,^67^ lending temporal support for this hypothesis.

We observed best benchmark concentrations below the current BLRV set by the CDC (3.5μg/dL or 0.17μM),^19^ with several genes affected by levels of Pb below this limit including *COL3A1*, *IGFBP5*, several transcription factors (*FOXD4L1* and *TWIST2*)^68,69^, and calcium binding factors *S100A11*^70^ and *GMP*^71^, suggesting current standards may not fully negate gene expression changes at low dose.

### Novel Pathways Affected by Environmentally Relevant Concentrations of Lead

Our final analysis evaluated which biological pathways Pb exposure can perturb. These results were exceptionally differentiation stage-specific, with the majority of effects on D18 (**Figure 7A**). The majority at any time point were associated with environmentally relevant concentrations of Pb (**Table 4**). Regarding GO pathway analysis, on D15, Pb-responsive genes had depressed expression related to DNA replication, cell cycling and cellular proliferation. We observed upregulation of pathways related to protein misfolding and cholesterol biosynthesis. Aberrations to cholesterol production are linked to cell cycle arrest,^72^ so on D15, Pb may be disrupting its biosynthesis, leading to a temporary hike in compensating factors and a depression of cell cycling. By D18, genes related to DNA regulation and damage were downregulated, as well as downregulation of genes involved in DNA replication and an increase in those involved Golgi vesicle formation. Golgi vesicle formation relies heavily on cholesterol, an essential component of the vesicle membrane.^73^ The disruptions in cholesterol biosynthesis and function suggests another by which Pb exposure affects cellular health. This is of particular relevance as aberrant production and metabolism of cholesterol is linked to neurodegenerative outcomes^74^ including risk factors such as the *APOE4* variant.^75^

**Table 4:** Median Benchmark Concentrations for Significantly Enriched Pathways Through the Gene Ontology Biological Process Database. Median BMCs were identified in BMDExpress using the Functional Classification tool. BMCs are presented here for each GO Biological Process pathway significantly enriched for Pb-responsive genes (p<0.05) in µM units across all analyzed timepoints. BMC: benchmark concentration.

| GO Biological Process Pathway Name | Day 6 | Day 9 | Day 12 | Day 15 | Day 18 |
| --- | --- | --- | --- | --- | --- |
| spindle assembly involved in female meiosis I | -- | -- | 1.87 | 0.09 | -- |
| cranial ganglion development | -- | 0.28 | -- | -- | -- |
| DNA topological change | -- | -- | -- | -- | 0.63 |
| regulation of DNA primase activity | -- | -- | -- | -- | 0.08 |
| positive regulation of DNA primase activity | -- | -- | -- | -- | 0.08 |
| nuclear cell cycle DNA replication initiation | -- | -- | -- | 0.19 | 1.31 |
| mitotic DNA replication initiation | -- | -- | -- | 0.19 | 1.31 |
| cell cycle DNA replication initiation | -- | -- | -- | 0.19 | 1.31 |
| positive regulation of mitotic cytokinesis | -- | -- | 2.47 | 0.137593 | 0.03 |
| ATF6-mediated unfolded protein response | -- | -- | -- | 0.39 | -- |
| positive regulation of chromosome condensation | -- | -- | 1.66 | -- | 1.58 |
| thymidine biosynthetic process | -- | 0.46 | -- | 0.31 | -- |
| pyrimidine deoxyribonucleoside biosynthetic process | -- | 0.46 | -- | 0.31 | -- |
| deoxyribonucleoside biosynthetic process | -- | 0.46 | -- | 0.31 | -- |
| positive regulation of mitochondrial ATP synthesis coupled electron transport | -- | -- | 2.83 | -- | -- |
| cysteine biosynthetic process | 1.1 | -- | -- | -- | -- |
| meiotic chromosome condensation | -- | -- | -- | -- | 0.27 |
| L-serine biosynthetic process | -- | -- | -- | -- | 1.57 |
| Golgi transport vesicle coating | -- | -- | -- | -- | 3.81 |
| COPI coating of Golgi vesicle | -- | -- | -- | -- | 3.81 |
| catecholamine secretion | -- | -- | -- | -- | 0.71 |
| positive regulation of meiotic cell cycle phase transition | -- | -- | -- | -- | 2.77 |
| protein maturation by [4Fe-4S] cluster transfer | -- | -- | 3.2 | -- | -- |
| apical junction assembly | -- | -- | 2.41 | -- | -- |
| positive regulation of long-chain fatty acid import into cell | -- | -- | -- | 1.29 | -- |
| dTMP biosynthetic process | -- | -- | -- | 0.29 | -- |
| sympathetic neuron projection guidance | -- | 0.46 | -- | -- | -- |
| regulation of isoprenoid metabolic process | -- | 0.4 | -- | -- | -- |
| regulation of ferroptosis | -- | -- | 1.63 | -- | -- |
| lncRNA-mediated post-transcriptional gene silencing | -- | -- | 1.69 | -- | -- |
| UDP metabolic process | -- | -- | -- | -- | 0.49 |
| replication-born double-strand break repair via sister chromatid exchange | -- | -- | -- | -- | 1.18 |
| regulation of RNA polymerase II transcription preinitiation complex assembly | -- | -- | -- | -- | 0.43 |
| pyrimidine ribonucleoside diphosphate metabolic process | -- | -- | -- | -- | 0.49 |
| positive regulation of protein localization to centrosome | -- | -- | -- | -- | 0.66 |
| positive regulation of dopamine metabolic process | -- | -- | -- | -- | 0.23 |
| positive regulation of catecholamine metabolic process | -- | -- | -- | -- | 0.23 |
| negative regulation of protein kinase activity by regulation of protein phosphorylation | -- | -- | -- | -- | 1.68 |
| intraciliary transport involved in cilium assembly | -- | -- | -- | -- | 0.002 |
| negative regulation of oxidative phosphorylation uncoupler activity | 0.19 | -- | -- | -- | -- |
| actin rod assembly | 3.01 | -- | -- | -- | -- |
| VEGF-activated platelet-derived growth factor receptor signaling pathway | -- | -- | 3.01 | -- | -- |
| regulation of telomerase RNA reverse transcriptase activity | -- | -- | 0.55 | -- | -- |
| regulation of Schwann cell differentiation | -- | -- | 2.63 | -- | -- |
| positive regulation of cell proliferation by VEGF-activated platelet derived growth factor receptor signaling pathway | -- | -- | 3.01 | -- | -- |
| regulation of heparan sulfate proteoglycan biosynthetic process | -- | -- | -- | 0.63 | -- |
| pyrimidine deoxyribonucleoside monophosphate biosynthetic process | -- | -- | -- | 0.29 | -- |
| positive regulation of exit from mitosis | -- | -- | -- | 0.04 | -- |
| negative regulation of mammary gland epithelial cell proliferation | -- | -- | -- | 0.7 | -- |
| leading strand elongation | -- | -- | -- | 0.07 | -- |
| depurination | -- | -- | -- | 0.51 | -- |
| chondroitin sulfate catabolic process | -- | -- | -- | 0.88 | -- |
| cholesterol biosynthetic process via lathosterol | -- | -- | -- | 0.04 | -- |
| cholesterol biosynthetic process via desmosterol | -- | -- | -- | 0.045 | -- |
| cellular response to cisplatin | -- | -- | -- | 0.26 | -- |
| aminergic neurotransmitter loading into synaptic vesicle | -- | -- | -- | 0.42 | -- |
| viral translational termination-reinitiation | -- | -- | -- | -- | 0.17 |
| protein retention in Golgi apparatus | -- | -- | -- | -- | 0.67 |
| positive regulation of melanocyte differentiation | -- | -- | -- | -- | 0.59 |
| positive regulation of DNA ligation | -- | -- | -- | -- | 0.34 |
| negative regulation of helicase activity | -- | -- | -- | -- | 0.46 |
| morphogenesis of a polarized epithelium | -- | -- | -- | -- | 2.2 |
| mitochondrial DNA repair | -- | -- | -- | -- | 0.54 |
| cellular response to heparin | -- | -- | -- | -- | 3.84 |

Our analysis of Pb-responsive genes with relation to MSigDB hallmarks and supplemental gene sets also revealed interesting trends. The p53 pathway, MYC targets, and MTORC1 were all significantly enriched at environmentally relevant concentrations. Given their role in cancer suppression,^76^ cellular proliferation,^77^ and cellular growth,^78^ respectively, this suggests Pb exposure may induce cancer-causing mechanisms within the differentiating cells, which has been suggested previously^79^ and this relationship may be relevant to neurodegenerative disease risk (**Figure 8**, **Table 5**).^80^

**Table 5:**
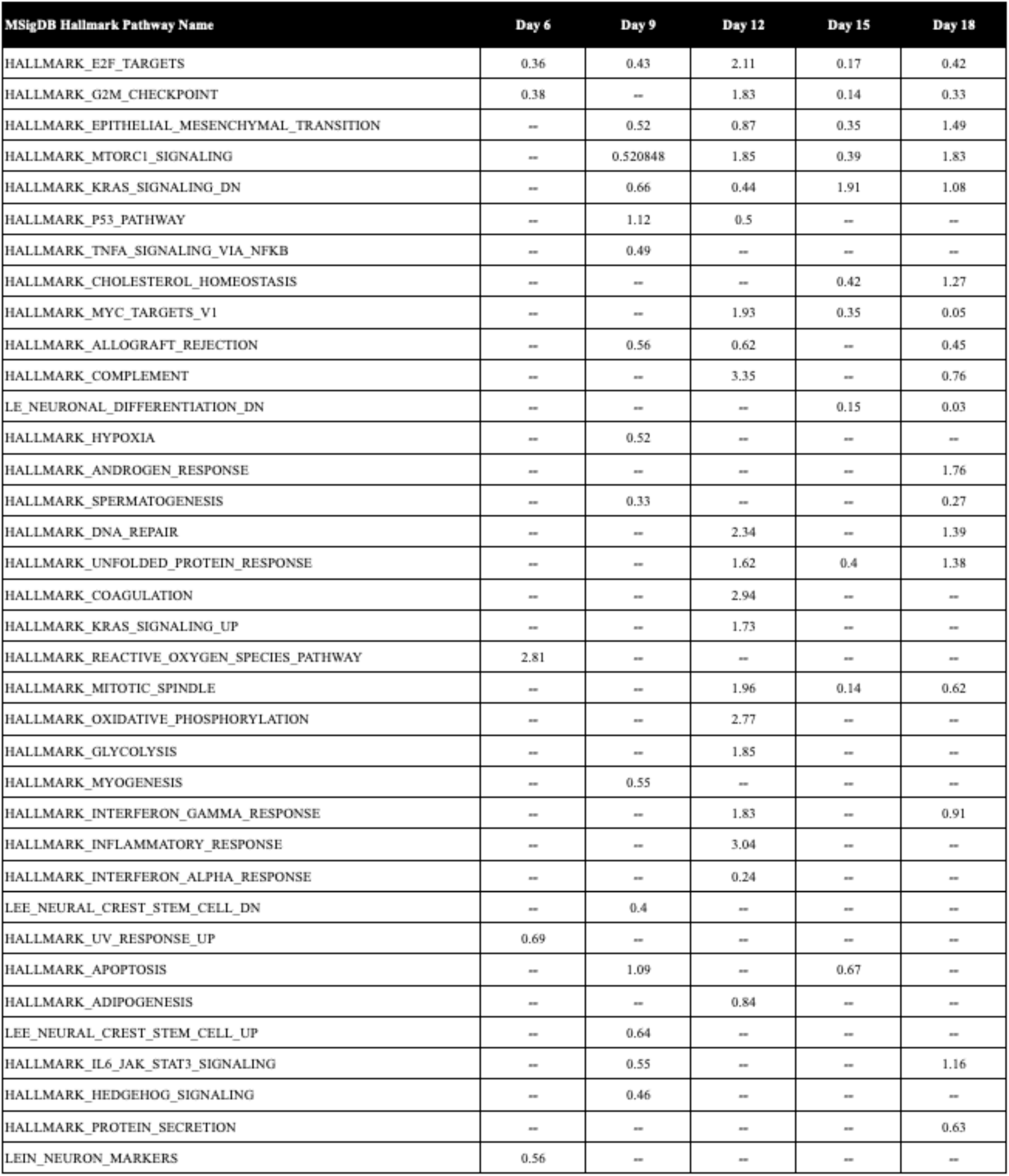
Median Benchmark Concentrations for Significantly Enriched Pathways through the Molecular Signatures Database. Median BMCs were identified in BMDExpress using the Functional Classification tool. BMCs are presented here for each MSigDB hallmark significantly enriched for Pb-responsive genes (p<0.05) in µM units across all analyzed timepoints. BMC: benchmark concentration

| MSigDB Hallmark Pathway Name | Day 6 | Day 9 | Day 12 | Day 15 | Day 18 |
| --- | --- | --- | --- | --- | --- |
| HALLMARK_E2F_TARGETS | 0.36 | 0.43 | 2.11 | 0.17 | 0.42 |
| HALLMARK_G2M_CHECKPOINT | 0.38 | -- | 1.83 | 0.14 | 0.33 |
| HALLMARK_EPITHELIAL_MESENCHYMAL_TRANSITION | -- | 0.52 | 0.87 | 0.35 | 1.49 |
| HALLMARK_MTORC1_SIGNALING | -- | 0.520848 | 1.85 | 0.39 | 1.83 |
| HALLMARK_KRAS_SIGNALING_DN | -- | 0.66 | 0.44 | 1.91 | 1.08 |
| HALLMARK_P53_PATHWAY | -- | 1.12 | 0.5 | -- | -- |
| HALLMARK_TNFA_SIGNALING_VIA_NFKB | -- | 0.49 | -- | -- | -- |
| HALLMARK_CHOLESTEROL_HOMEOSTASIS | -- | -- | -- | 0.42 | 1.27 |
| HALLMARK_MYC_TARGETS_V1 | -- | -- | 1.93 | 0.35 | 0.05 |
| HALLMARK_ALLOGRAFT_REJECTION | -- | 0.56 | 0.62 | -- | 0.45 |
| HALLMARK_COMPLEMENT | -- | -- | 3.35 | -- | 0.76 |
| LE_NEURONAL_DIFFERENTIATION_DN | -- | -- | -- | 0.15 | 0.03 |
| HALLMARK_HYPOXIA | -- | 0.52 | -- | -- | -- |
| HALLMARK_ANDROGEN_RESPONSE | -- | -- | -- | -- | 1.76 |
| HALLMARK_SPERMATOGENESIS | -- | 0.33 | -- | -- | 0.27 |
| HALLMARK_DNA_REPAIR | -- | -- | 2.34 | -- | 1.39 |
| HALLMARK_UNFOLDED_PROTEIN_RESPONSE | -- | -- | 1.62 | 0.4 | 1.38 |
| HALLMARK_COAGULATION | -- | -- | 2.94 | -- | -- |
| HALLMARK_KRAS_SIGNALING_UP | -- | -- | 1.73 | -- | -- |
| HALLMARK_REACTIVE_OXYGEN_SPECIES_PATHWAY | 2.81 | -- | -- | -- | -- |
| HALLMARK_MITOTIC_SPINDLE | -- | -- | 1.96 | 0.14 | 0.62 |
| HALLMARK_OXIDATIVE_PHOSPHORYLATION | -- | -- | 2.77 | -- | -- |
| HALLMARK_GLYCOLYSIS | -- | -- | 1.85 | -- | -- |
| HALLMARK_MYOGENESIS | -- | 0.55 | -- | -- | -- |
| HALLMARK_INTERFERON_GAMMA_RESPONSE | -- | -- | 1.83 | -- | 0.91 |
| HALLMARK_INFLAMMATORY_RESPONSE | -- | -- | 3.04 | -- | -- |
| HALLMARK_INTERFERON_ALPHA_RESPONSE | -- | -- | 0.24 | -- | -- |
| LEE_NEURAL_CREST_STEM_CELL_DN | -- | 0.4 | -- | -- | -- |
| HALLMARK_UV_RESPONSE_UP | 0.69 | -- | -- | -- | -- |
| HALLMARK_APOPTOSIS | -- | 1.09 | -- | 0.67 | -- |
| HALLMARK_ADIPOGENESIS | -- | -- | 0.84 | -- | -- |
| LEE_NEURAL_CREST_STEM_CELL_UP | -- | 0.64 | -- | -- | -- |
| HALLMARK_IL6_JAK_STAT3_SIGNALING | -- | 0.55 | -- | -- | 1.16 |
| HALLMARK_HEDGEHOG_SIGNALING | -- | 0.46 | -- | -- | -- |
| HALLMARK_PROTEIN_SECRETION | -- | -- | -- | -- | 0.63 |
| LEIN_NEURON_MARKERS | 0.56 | -- | -- | -- | -- |

### Limitations

SH-SY5Y cells are an imperfect model of neural differentiation as the *in vitro* differentiation into a single cell type does not recapitulate the dynamic context of the brain during periods of neurodevelopment. Future work could use pluripotent stem cells and experimental methods to differentiate these into multiple cell types to more closely replicate this process. SH-SY5Y are also cancerous in origin, being derived from a neuroblastoma. This influences their differentiation along with other properties including viability and metabolism^81^, and the use of a truly pluripotent stem cell line would address this limitation as well.

### Summary and Public Health Significance

Our data demonstrates that continuous Pb exposure during neural differentiation has significant impacts on gene expression, many of which are differentiation-stage specific and present at relatively low exposure levels. Exposure was associated with the differential expression of genes involved in cell cycling and DNA replication and repair, protein misfolding, and cholesterol synthesis, all of which are implicated in neurodegenerative disease risk. These results support the hypothesis that environmental toxicants such as Pb perturb proper neuronal development and contribute to the risk of neurodegenerative disease risk later in life.

### Acronyms

Pb (lead), RNA (ribonucleic acid), DNA (deoxyribonucleic acid), BLL (blood lead level), BLRV (blood lead reference value), CDC (Centers for Disease Control and Prevention), RA (retinoic acid), UM (University of Michigan), FDR (false discovery rate), FC (fold change), BMC (benchmark concentration), GO (Gene Ontology), MSigDB (Molecular Signatures Database)

## Funding

This work was supported by funding from the following sources: National Institute of Environmental Health Sciences (NIEHS) Grants R35 (ES031686), R01 (ES028802), and the Michigan Lifestage Environmental Exposures and Disease (M-LEEaD) NIEHS Core Center (P30 ES017885), Institutional Training Grant T32 (ES007062), Institutional Training Grant T32 (HD079342), and National Institute on Aging (NIA) Grant R01 (AG072396).

## Disclosures

Dr. Dolinoy reports serving as an expert legal consultant for toxicology and epigenetics lawsuits, providing expert testimony for various legal cases. None of these disclosures played a role in the analysis, interpretation, decision to publish, or preparation of this manuscript. None of the other authors has any disclosures to make.

