## Supplementary Table 1 for "Environmentally Relevant Lead Exposure Impacts Gene Expression in SH-SY5Y Cells Throughout Neuronal Differentiation"

| SH-SY5Y Differentiation Media Description | | |
|---|---|---|
| **Basic Growth Media** | | |
| **Component** | **Volume for 50mL** | **Final Concentration** |
| EMEM | 41.5mL | |
| hiFBS | 7.5mL | 15% |
| 100x Pen/Strep | 0.5mL | 1x |
| 200mM Glutamine | 0.5mL | 2mM |
| **Differentiation Media #1** | | |
| **Component** | **Volume for 50mL** | **Final Concentration** |
| EMEM | 48mL | |
| hiFBS | 1.3mL | 2.5% |
| 100x Pen/Strep | 0.5mL | 1x |
| 200mM Glutamine | 0.5mL | 2mM |
| 5mM RA | 0.1mL | 10µM |
| **Differentiation Media #2** | | |
| **Component** | **Volume for 50mL** | **Final Concentration** |
| EMEM | 49mL | |
| hiFBS | 0.5mL | 1% |
| 100x Pen/Strep | 0.5mL | 1x |
| 200mM Glutamine | 0.5mL | 2mM |
| 5mM RA | 0.1mL | 10µM |
| **Differentiation Media #3** | | |
| **Component** | **Volume for 50mL** | **Final Concentration** |
| Neurobasal | 47mL | |
| 100x B-27 | 1mL | |
| 1M KCl | 1mL | |
| 100x Pen/Strep | 0.5mL | |
| 200mM Glutamax-I | 0.5mL | 2mM |
| 10µg/mL BDNF | 0.25mL | 50ng/mL |
| 1M db-cAMP | 0.1mL | 2mM |
| 5mM RA | 0.1mL | 10µM |

**Table S1: Description of Media Conditions for SH-SY5Y Differentiation.** Adapted from Shipley, et al., 2016.
