## Supplementary Table 4 for "Environmentally Relevant Lead Exposure Impacts Gene Expression in SH-SY5Y Cells Throughout Neuronal Differentiation"

| | Day 9 v. Day 6 | Day 12 v. Day 6 | Day 15 v. Day 6 | Day 18 v. Day 6 |
|---|---|---|---|---|
| Up | 63 | 1437 | 1743 | 2377 |
| Down | 35 | 723 | 864 | 1612 |
| Total Unique DEGs | 98 | 2160 | 2598 | 3989 |

**Table S4: Summary of Differentially Expressed Gene Counts.** SH-SY5Y cells, baseline changes in gene expression with no exposure. DEG: differentially expressed genes.
