## Supplementary Figure 1 for "Environmentally Relevant Lead Exposure Impacts Gene Expression in SH-SY5Y Cells Throughout Neuronal Differentiation"

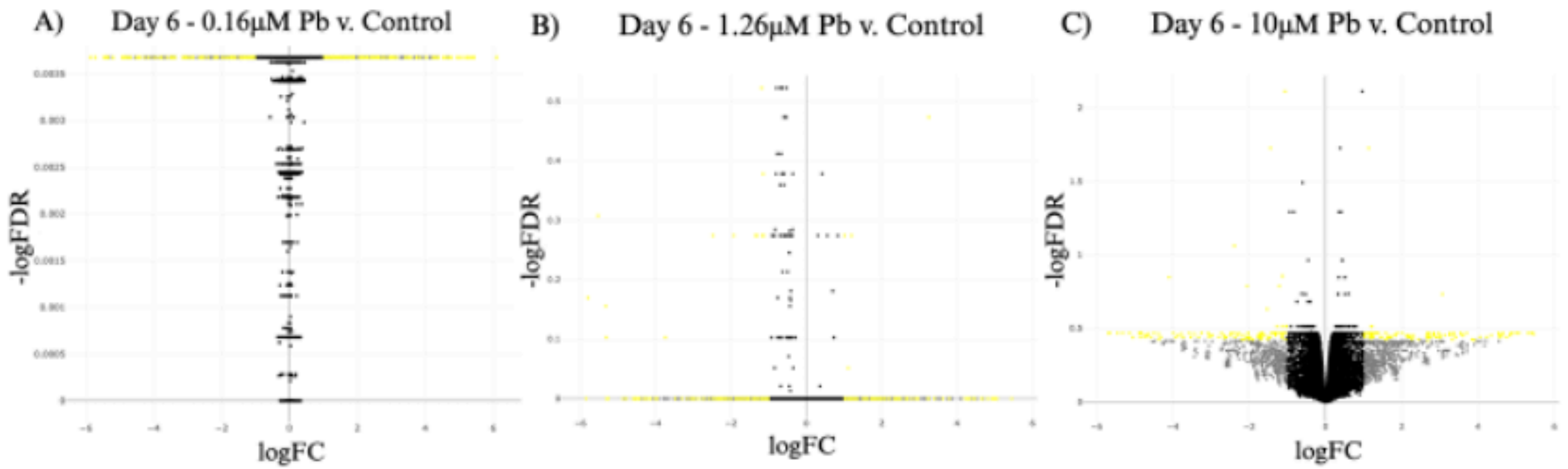


**Figure S1: Differential Gene Expression Associated with Lead Exposure During SH-SY5Y Differentiation – Day 6.** Differential gene expression during differentiation was calculated for each dose, 0.16μM Pb (A), 1.26μM Pb (B), 10μM (C), relative to control cells Significantly differentially expressed genes ($|log_2FC| > 1$, FDR < 0.05, p-value < 0.05) denoted in red, differentially expressed genes ($|log_2FC| > 1$, p-value < 0.05) and significant genes (FDR < 0.05 and p-value < 0.05) denoted in yellow and blue, respectively. Genes with insignificant changes in expression ($|log_2FC| > 1$, p-value and FDR > 0.05) denoted in grey and genes with no detectable change in expression between conditions denoted in black. Corresponding counts of differentially expressed genes at each time point can be found in **Table S2**.
