## Supplementary Figure 2 for "Environmentally Relevant Lead Exposure Impacts Gene Expression in SH-SY5Y Cells Throughout Neuronal Differentiation"

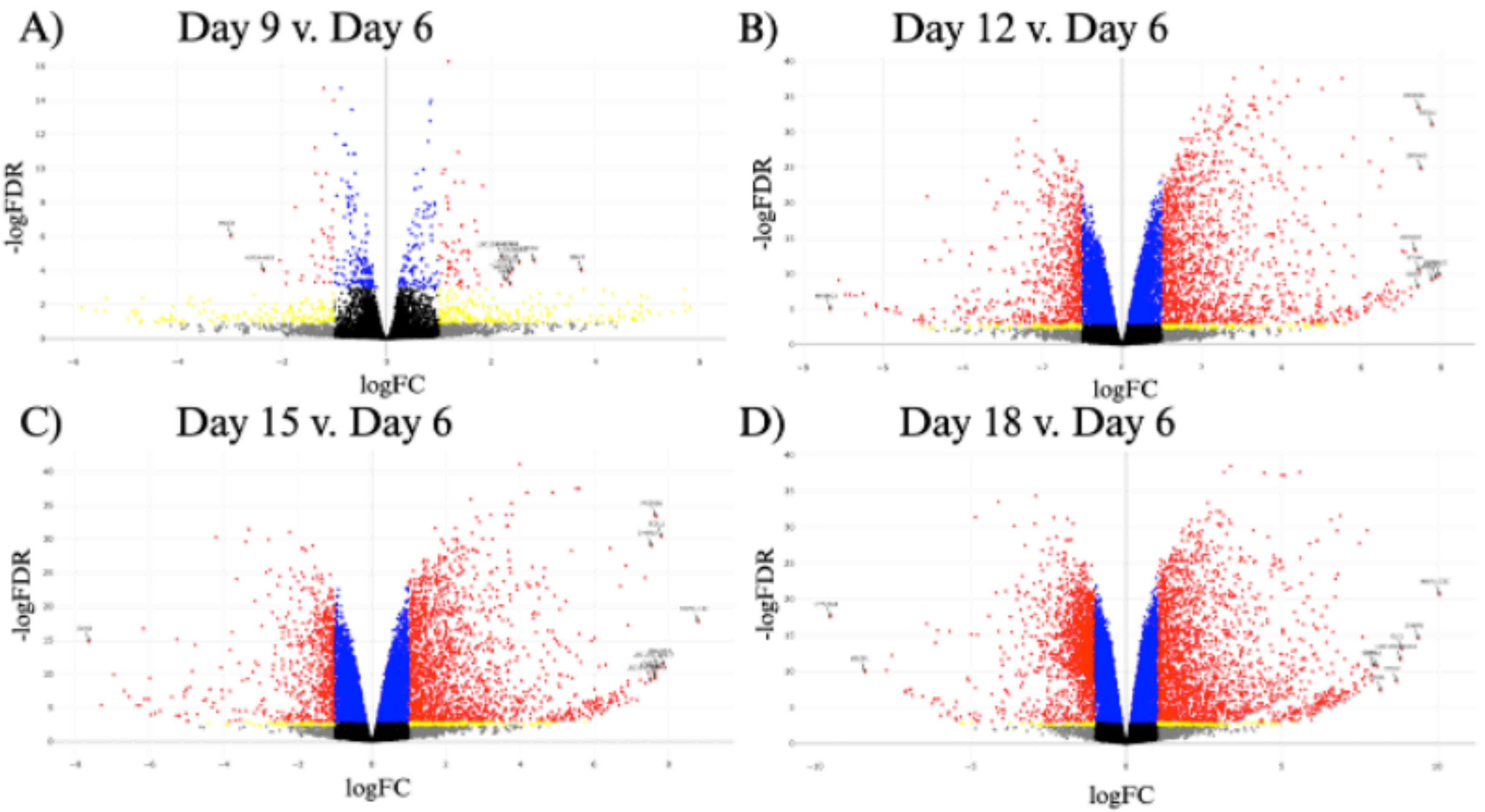


**Figure S2: Changes in Gene Expression During SH-SY5Y Differentiation in Control (Unexposed) Samples.** SH-SY5Y cells were differentiated according to previously published protocols (Shipley et al., 2019) and RNA was extracted every three days. Differential gene expression during differentiation was calculated relative to the first time point (i.e., Day 6).
