## Supplementary Figure 3 for "Environmentally Relevant Lead Exposure Impacts Gene Expression in SH-SY5Y Cells Throughout Neuronal Differentiation"

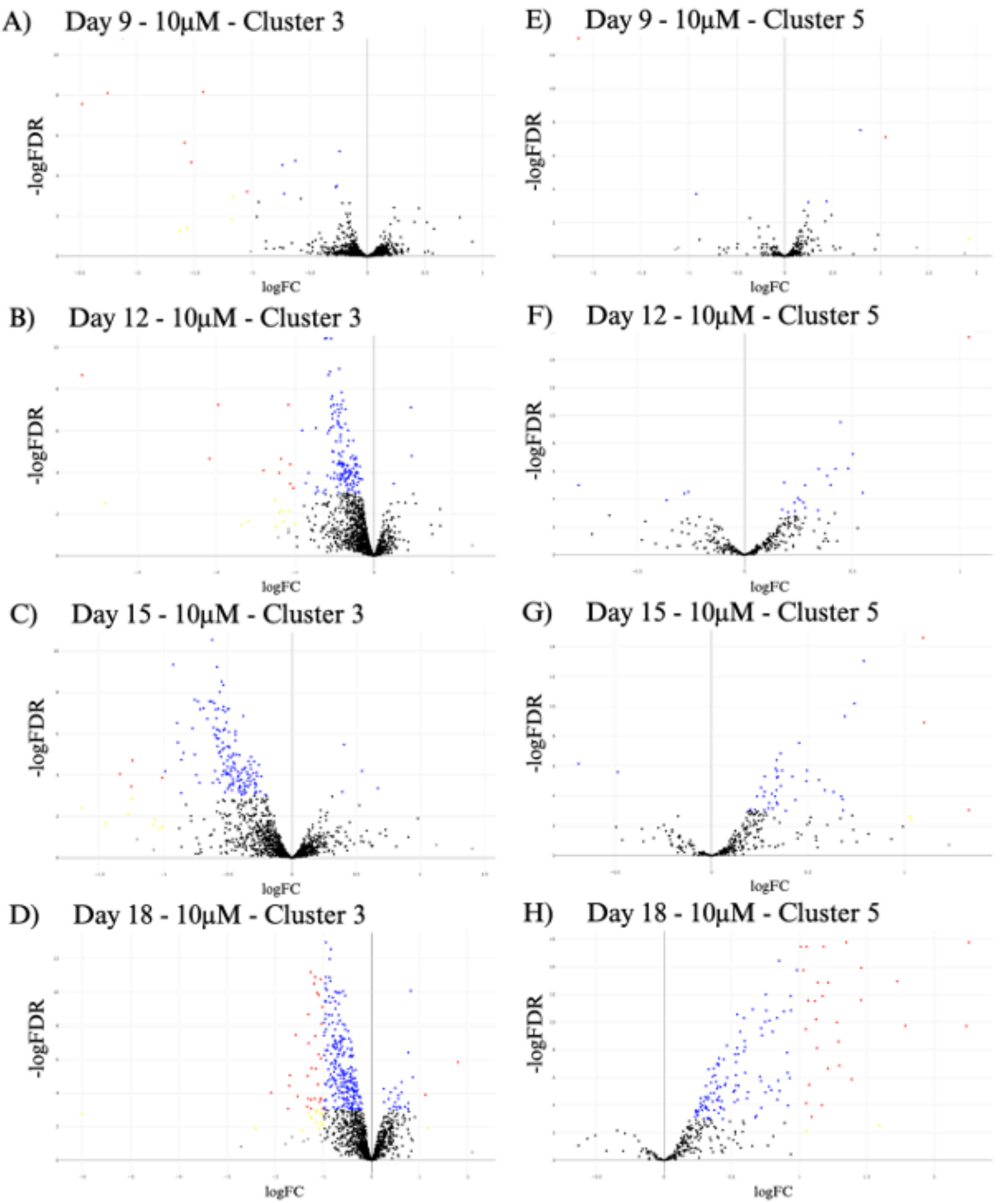


**Figure S3: Differential Expression of Genes in Clusters 3 and 5 Associated with Lead Exposure.** Significantly differentially expressed genes ($|\log_2 FC| > 1$, FDR < 0.05, p-value < 0.05) denoted in red, differentially expressed genes ($|\log_2 FC| > 1$, p-value < 0.05) and significant genes (FDR < 0.05 and p-value < 0.05) denoted in yellow and blue, respectively. Genes with insignificant changes in expression ($|\log_2 FC| > 1$, p-value and FDR > 0.05) denoted in grey and genes with no detectable change in expression between conditions denoted in black.
